# Impact of maternal iron deficiency anaemia on fetal iron status and placental iron transporters in human pregnancy

**DOI:** 10.1101/2022.06.29.498050

**Authors:** Sreenithi Santhakumar, Rekha Athiyarath, Anne George Cherian, Vinod Joseph Abraham, Biju George, Paweł Lipiński, Eunice Sindhuvi Edison

**Affiliations:** Department of Haematology, Christian Medical College, Vellore, India; Department of Community Health and Development, Christian Medical College, Vellore, India; Institute of Genetics and Animal Biotechnology, Polish Academy of Sciences, Jastrzębiec, ul. Postępu 36A, 05-552 Magdalenka Poland

**Keywords:** Pregnancy, Placenta, Iron deficiency anemia, Iron regulators

## Abstract

Iron deficiency anaemia is associated with maternal morbidity and poor pregnancy outcomes. Placenta expresses both haem and non-haem iron transport proteins. The aim of the study is to examine the expression of placental iron trafficking molecules and associate them with maternal and neonatal iron status. Pregnant women who received prenatal care at the department of community health and development, Christian Medical College, Vellore, India for childbirth were recruited between 2016-2018. Pregnant women who were 18-35 years old with gestational age (GA) of ≥36weeks were eligible to participate in the study. In a prospective cohort of pregnant women, 22% were iron deficiency anaemia (IDA) and 42% were iron replete. Pregnant women in the different groups were mutually exclusive. Samples were collected (Maternal blood, placental tissue, and cord blood) from pregnant women with gestational age of ≥38 weeks at the time of delivery. Mean gestational age at first visit and delivery was 12.8±2.72 weeks and 39±1.65 weeks, respectively. Hemoglobin (9.3±0.9g/dl) and ferritin (15.4(0.8-28.3) ng/ml) levels at delivery were significantly decreased in IDA as compared to other groups. The foetal haemoglobin and ferritin levels were in the normal range in all groups. We further analysed the expression of iron transport genes in the placenta in the iron replete controls and the IDA group. Under maternal iron insufficiency, the expression of placental iron transporters DMT1 and FPN1 were upregulated at the transcriptional level. There was no correlation of maternal and cord blood hepcidin with foetal iron status in IDA. Thus, placental iron traffickers respond to maternal iron deficiency by increasing their expression and allowing sufficient iron to pass to the foetus.

## 1 Introduction

Iron deficiency is a well-known micro nutritional deficiency causing severe anaemia in maternal women and increases the risk of neuronal impairment in neonates^1^. Iron deficiency anaemia (IDA) affects 1.7 billion people globally; among them pregnant women are the most vulnerable population^2^. In India, iron deficiency is most common cause of anaemia in 58% of pregnant women^3^. During pregnancy, a net cost of 1000mg of iron is required for the developing fetal-placental unit and increased maternal erythrocyte mass expansion, of which, one third of iron is utilised for establishing adequate iron stores in neonates at birth^4^. Placenta dynamically transports maternal iron via syncytiotrophoblasts towards the fetus and balances iron levels between mother and the fetus. Studies have shown the localisation of various iron transporters in placental microvillus and basal membranes, but their relevant mechanisms are poorly understood. The influence of maternal iron status towards the regulation of placental iron transport and fetal supply are less explored.

Hepcidin, the systemic regulator of iron bioavailability, decreases as pregnancy progresses and reaches to undetectable levels at the end of third trimester^5^. Maternal hepcidin regulates iron absorption towards fetal iron transport and fetal derived hepcidin regulates placental iron transporters and determines rate of iron transfer to fetus^6^. The maternal hepcidin contribution towards placental iron transfer was noted in an isotope study, where pregnant women (ingested with ^57^FeSO_4_) with undetectable level of serum hepcidin had increased radioisotope transfer to their fetus in comparison to detectable levels of serum hepcidin^7^. Increased fetal hepcidin levels reported in transgenic mice overexpressing hepcidin was able to regulate placental ferroportin and leading to severe iron deficiency and lethal^8^.

Growth Differentiation Factor 15 (GDF15), a TGFβ family member known to be involved in embryonic development, significantly increases during pregnancy. GDF15 has shown to be expressed strongly in placenta but function is unknown. Data from secondary iron overload states such as in β-thalassaemia and congenital dyserythropoietic anaemia (CDA) shows that GDF15 suppresses hepcidin leading to regulation of iron absorption^9^.

Placental iron transporters including heme and non-heme iron transporters are in syncytiotrophoblasts, whose interplay in placental iron acquisition has to be studied further^10^. Bradley and co-workers analysed 22 pregnant women placental tissues at different gestational ages^11^. They demonstrated that Iron regulatory protein isoforms IRP1 and IRP2 activity is present throughout gestation and responds to foetal iron status. IRP1 activity was the mainstay for post transcriptional regulation of ferritin (FT) and ferroportin (FPN) in placenta^11^. Chong’s immunohistochemical study exhibited isoforms of dimetal transporters (DMT1) such as DMT1A containing IRE in its 3’ UTR and DMT1B without IRE were expressed in syncytiotrophoblasts, were responsible for cellular iron transport in placenta. ^12^ Recent study using IRP1 knockout iron deficient mice illustrated that placental iron regulators FPN and transferrin receptor(TFRC) function is regulated by IRP1 activity in response to maternal iron deficiency ^13^. Most of the studies have explored IRP1 involvement in placental iron regulation, but the mechanism of placental IRP2 either in normal or iron deficient condition remains to be characterised.

Here we investigated the changes of hepcidin, ferritin, GDF15 and haematological parameters in iron deficient pregnant women and compared them to iron replete pregnant women. We also compared maternal and fetal iron status with mRNA expression and protein levels of placental iron transporters.

## 2 Materials and Methods

### 2.1 Study population

The study was approved by Institutional Review Board (Ethics committee) of Christian Medical College (CMC) at Vellore, India, and informed written consent was obtained from all the study participants. This is a cross sectional study conducted in pregnant women between the year 2016-2018. Subjects who visited the antenatal clinic at department of Community Health and Development, Christian Medical College, Vellore for childbirth were screened and subjects who fulfilled the inclusion criteria were included in the study (Age-18-35 years and a gestational age (GA) of ≥36weeks). Pregnant women with gestational diabetes, pregnancy induced hypertension (PIH), hypothyroidism, previous caesarean section, bacterial or viral infections during onset of labour, twin pregnancy, who received transfusion during delivery were not included in the study. Daily oral iron supplementation with 60 mg of elemental iron was recommended for all pregnant women visited our antenatal clinic. A detailed proforma was recorded including type of delivery, placenta size and newborn details such as sex, baby weight.

Maternal blood samples were collected at admission (GA≥38 weeks) prior to or immediately after delivery. During delivery, the umbilical cord was clamped, cut and cord blood was collected. Placental tissue was obtained and processed within an hour of delivery.

For the analysis, Iron deficiency anaemia in pregnancy (IDA) was defined as Hb level of <10.5g/dl with a ferritin level <30ng/ml ; iron replete subjects (control) Hb>10.5g/dl and ferritin >30ng/ml.

### 2.2 Haematological and biochemical Assessment

Complete blood counts (CBC) were carried out on maternal peripheral blood and cord blood samples using an automated haematology analyser (Sysmex KX21). Serum ferritin was analysed using a chemiluminescence immunoassay using the Advia Centaur, Siemens XPI. Serum hepcidin was quantified by using an enzyme immunoassay method from DRG, GmbH according to the manufacturer’s protocol. GDF15 was quantified in serum using an ELISA method (R&D Systems, Inc. MN, USA).

### 2.3 Placenta collection and processing

Placenta collected during delivery was processed within 1 hour following delivery. Amniotic membranes were removed from the placenta and tissue of 0.5-0.8cm thickness was incised from red cotyledons and below the amniotic membrane side of the placenta and a deep cut was avoided. The dissected tissues were stored in RNAlater (Ambion) at -80□C until analysis.

We used variable number tandem repeat (VNTR) analysis using five markers to rule out maternal contamination in the placental tissues. Briefly DNA was extracted from maternal peripheral blood, cord blood and placental tissues. A multiplex PCR for five short tandem repeat (STR) markers (*ACTBP2, FES, THO1, VWF* and *F13A1*) was carried out using fluorescently labelled primers followed by capillary electrophoresis. It was confirmed that all placental tissue samples collected had fetal origin (Supplemental Figure 1).

### 2.4 RNA extraction and PCR Arrays

Total RNA was extracted from the frozen placental tissue using the Protein and RNA Isolation (PARIS) kit (Qiagen) following the manufacturer’s instructions. RNA was reverse transcribed and converted into cDNA using RT^2^ first strand Kit (QIAGEN). The cDNA was then diluted with nuclease-free water and added to the RT^2^ qPCR SYBR green Master Mix (SA Biosciences, Frederick MD). 25μl of the experimental cocktail was added to each well of the custom PCR array (SA Biosciences, Frederick MD) (Supplemental Table 1). Real-Time PCR was performed on the 7500 QPCR System (Applied Biosystems model) and used SYBR green detection. All data from the PCR was analysed by SA Bioscience’s PCR array data analysis web portal. Plate-to-plate variation was controlled by normalizing gene expression to β-actin and control placenta by using the 2^-ΔΔCt^ method.

### 2.5 Protein expression of placental Fe transporters by immunoblotting

Placental tissues were lysed by homogenization in cell disruption buffer (PARIS kit, Ambion) according to the manufacturer’s protocol. Protein concentration was quantified using Bradford assay. All samples were prepared in Lamaelli buffer with reducing agent β-mercaptoethanol. 50µg of samples used for FPN1 were not pre heated. For DMT1, samples were prepared in Lamaelli buffer without reducing agent and was not pre-heated. For all other proteins, 30µg of samples were boiled at 100°C for 5 mins. Protein size markers (Bio-Rad precision plus protein standards) was loaded without heating. Tissue lysates were separated by SDS-PAGE gels (4%-12%) and transferred to polyvinylidene difluoride fluorescence membranes (Millipore, Billerica, MA, USA). Non-fat dry milk (NFDM-10%) was used to block the membranes and probed with primary antibody diluted in 5% NFDM diluted in TBS buffer with 0.1% Tween20 and kept at 4°C for overnight. Membranes were rinsed and probed with secondary antibody for 1.5hr in NFDM blocking buffer containing 0.1% Tween20. The primary and secondary antibodies are listed in Supplemental Table 2. The bands were visualised using chemiluminescence ECL system (Super signal west femto, Thermo Scientific). The protein bands were detected by FluorChem E system using digital darkroom software. Band intensities were quantified by densitometric analysis using ImageJ software.

### 2.6 Alternative transcripts of placental iron traffickers

We selected eighteen iron metabolising genes involved in placental iron homeostasis and their alternative transcript data were retrieved from Ensembl website. Genes include DMT1, TFRC, FPN1, STEAP3, SLC46A1, HIF1A, ACO1, IREB2, GDF15, TWSG1, SP1, TP53, GAPDH, HFE, CD163, LRP1, FLVCR1, PGF. Forty-six primer sets were designed to specifically amplify the main and alternative transcripts of these genes. Of these 46 primer sets, 23 transcripts were found to be expressed in the placental tissue by qualitative PCR. Selective amplification of these transcripts was qualitatively confirmed using two controls and IDA samples. Quantitative PCR was performed for 15 transcripts of eight genes. Relative quantification was done by using 2^**-**ΔΔ**Ct**^ method.

### 2.7 Statistical Analysis

Statistical analysis of the data was carried out using the software SPSS, version 20. For categorical data, the Chi-square test was used. Appropriate statistical tests including t test for continuous variables, analysis of variance (ANOVA) for comparison of groups, Mann– Whitney, and Kruskal–Wallis for nonparametric data were used. Associations between fetal parameters with maternal and placental factors were evaluated using univariate linear regression.

## 3 Results

### 3.1 Subject characteristics

In this prospective study we enrolled 138 pregnant women who visited the antenatal clinic. Fourteen subjects were excluded due to the unavailability of either maternal or cord blood serum samples(Supplemental Figure 2). All had received iron supplements (60mg elemental Fe/day till delivery) irrespective of the hemoglobin levels at first visit. Of the 124 pregnant women, 47% were primigravida, 44% second gravida and 9% multigravida. Mean gestational age at first visit and delivery was 12.8±2.72 weeks and 39±1.65 weeks, respectively. Six percent of pregnant subjects delivered preterm (<37weeks of gestation), 45% delivered early term (37-39 weeks), 42.8% delivered at full term (39-42 weeks), 7% delivered late term (41-42weeks) and one person delivered post term(≥42weeks). With respect to fetal gender, sixty-nine were males and fifty-five were females.

Primary aim of the study was to understand placental iron transport in pregnant women with iron deficiency anemia and healthy controls. We recruited pregnant women based on hemoglobin levels at the time of admission to labor ward. We classified groups as IDA and healthy controls based on the hemoglobin and ferritin levels at delivery(Figure 1).

**Fig 1:**
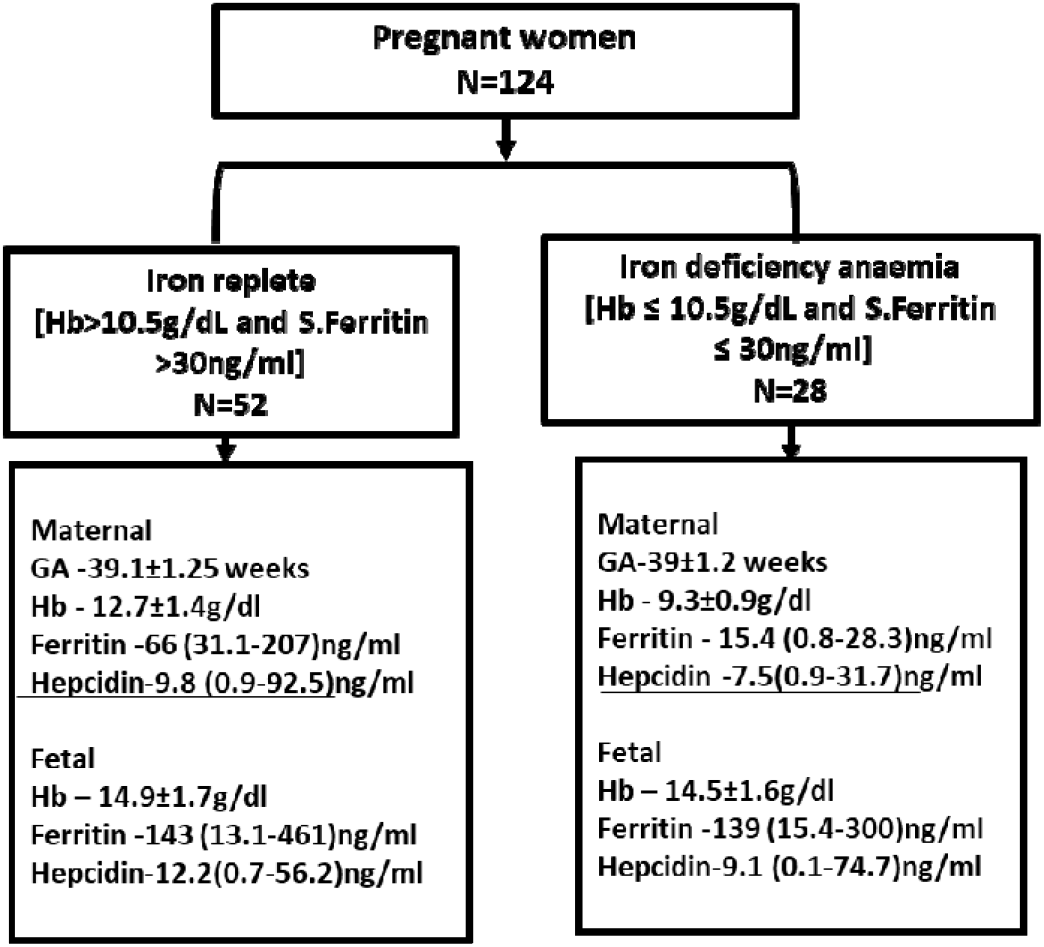
Groups classified based on Hemoglobin and ferritin levels. Listed gestational age(GA), Hemoglobin (Hb),serum ferritin and hepcidin levels of maternal and cord blood of each group.

Based on the inclusion criteria, among 124 subjects, 28 subjects had IDA (Hb≤10.5g/dL and ferritin values ≤30ng/L) and 52 were iron replete (Hb>10.5g/dL and ferritin >30ng/L). Remaining subjects failing to meet the inclusion criteria were excluded from further analyses.

The maternal mean age in the IDA was 24±3 and 25±3 years in the controls. At delivery, mean gestational age of IDA was 273±9 days and 274±8.8 days in controls, respectively. The mean birth weight of term neonates was 2.95±0.34 kg in IDA and 3±0.44 kg in controls.

Most (34/50;68%) of the subjects were classified to have mild anemia. In comparison to haemoglobin levels at the first antenatal visit, there was significant decrease in haemoglobin levels at delivery in IDA (p=0.000) and increase in control group at delivery (p=0.000) (Table.1).

The mean Hb concentration in term neonates in IDA was 14.5±1.6 g/dl and MCV 81.4±9.3 fL. The cord serum ferritin was 139 (15.4-300) ng/ml in the IDA group, (n=28). The demographic and laboratory parameters of the mother and their fetuses are presented in Table 1.

**Table 1:**
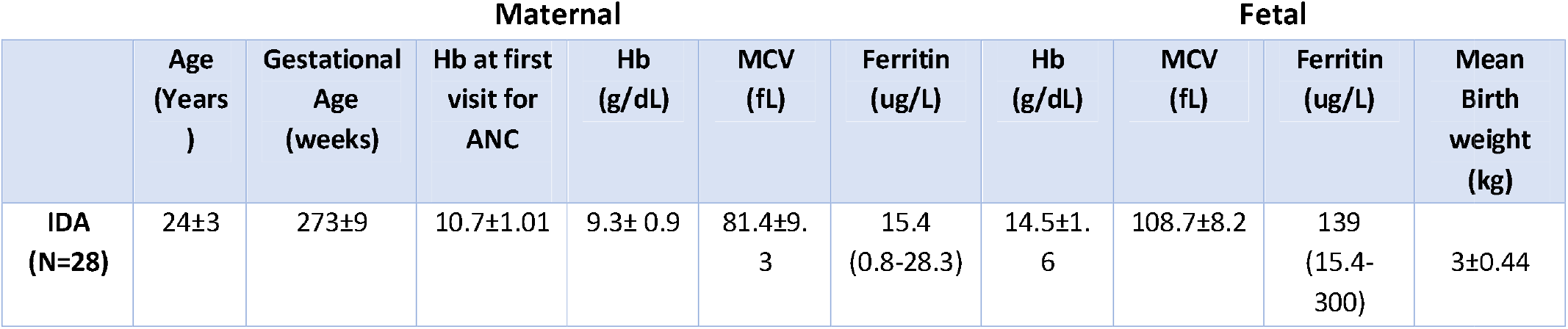
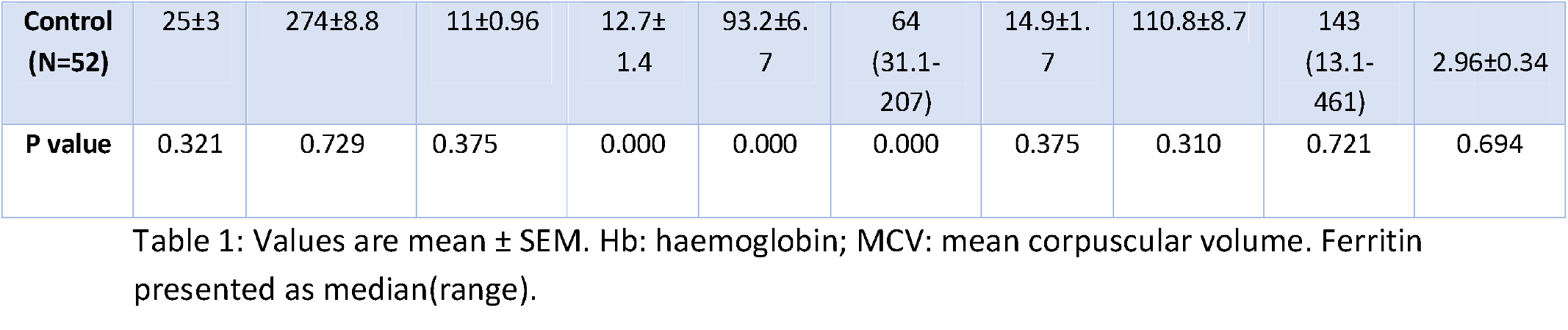
Baseline parameters of study groups

### 3.2 Assessment of iron status indicators in normal and iron deficient anaemia in pregnancy

The median level of maternal hepcidin was found to be 6.9 (0.9-19.5) ng/ml in IDA and 7.6 (0.9-19.3) ng/ml in controls (p=0.512). The median level of fetal hepcidin was 9.1 (0.2-74.7) ng/ml and 11.6 (0.9-54.8) ng/ml in the IDA and controls, respectively(p=0.686). Increased GDF15 levels was found in both groups with a median of 36040 (11910-66255) pg/ml in iron deficient mothers and 31070 (11477-67330) pg/ml in control group(p=0.365). Normal levels of GDF15 were observed in the cord blood of both IDA and controls [3840 (1880-7187) pg/ml and 3957 (2435-8542) pg/ml respectively (p=0.396) (Figure 2).

**Fig 2:**
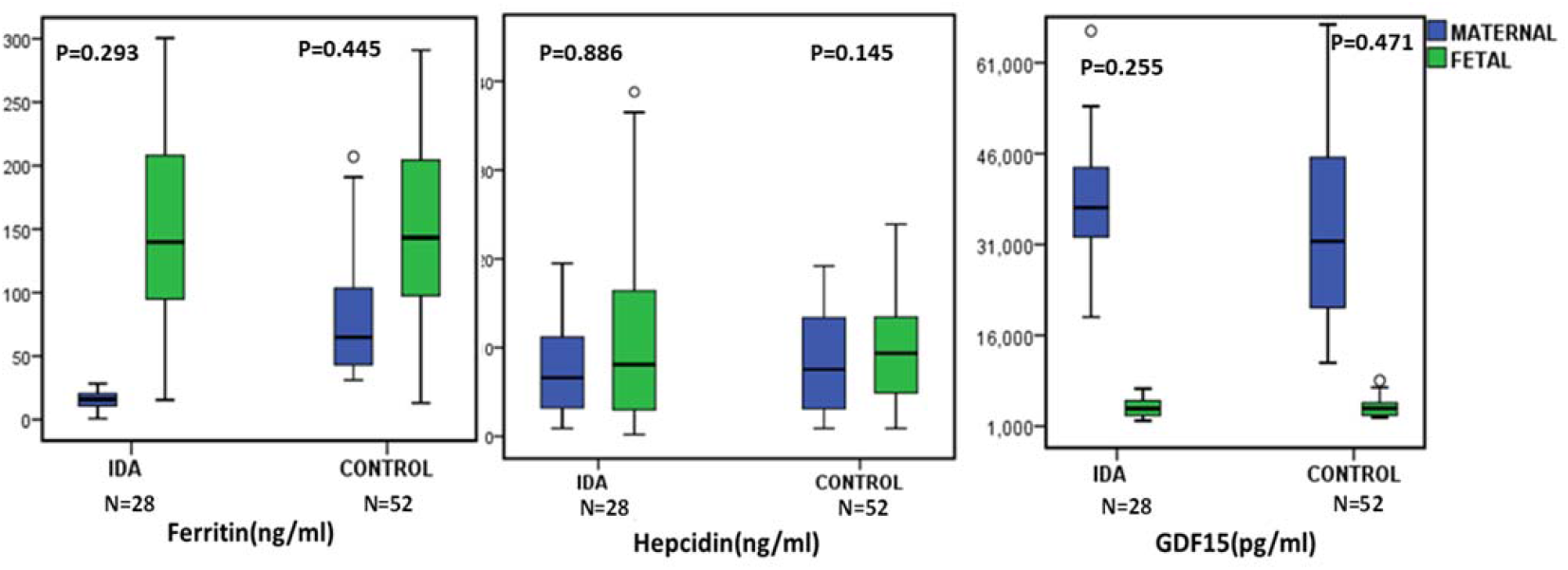
Ferritin, hepcidin and GDF15 levels in maternal and fetal cord blood serum. Fig 2: Ferritin, hepcidin and GDF15 levels in maternal and fetal cord blood serum compared between IDA and control group. The data are presented as mean±SD. Statistical differences between groups was determined by Mann-Whitney U rank-sum test for non-normally distributed values.

### 3.3 Association of maternal and fetal iron status indicators

In IDA, fetal cord blood Hb and ferritin levels were significantly higher than maternal Hb and serum ferritin respectively (p=0.000; p=0.000) (Table 1). Serum ferritin had positive correlation with maternal Hb levels in IDA (r=0.421, p=0.026). Hepcidin and ferritin levels were independent of each other in both maternal and cord blood.

By using univariate regression analysis, we found that fetal hemoglobin was associated with fetal ferritin level(β=-0.360;P<0.05). Maternal hepcidin: ferritin ratio was significantly higher in IDA (p=0.000) than controls. However, there was no association between maternal and cord blood hepcidin: ferritin ratio. Conversely, in controls, increased maternal hepcidin was associated with increased fetal hepcidin and fetal hepcidin: ferritin ratio respectively (r=0.442, p=0.001; r=0.379, p=0.006). Association between fetal ferritin and fetal hepcidin showed trend towards significance(r=0.273,p=0.052). Logarithmic fetal hepcidin was related to maternal ferritin (β=0.385 ;P=0.047). Fetal GDF15 was related to fetal hepcidin-ferritin ratio (β=0.476 ;P=0.014). Interestingly, multigravida pregnant women had significantly lower maternal hepcidin levels as compared to primigravida (p=0.014).

In both the groups, GDF15 was significantly higher in maternal serum as compared to cord blood levels (p=0.000). GDF15 did not influence hepcidin and ferritin levels in both mother and fetus. Interestingly, we observed that maternal GDF15 had negative association with fetal hepcidin: ferritin ratio in IDA (r= -0.439; p=0.025).

### 3.4 Analysis of Placental iron transporters and regulators

The expression of iron metabolising genes in maternal and fetal iron transfer were analysed at mRNA and protein level. Of the six differentially expressed genes, iron transporters (*DMT1, FPN1*), cellular iron regulator *IREB2*, known hepcidin suppressor *GDF15* and its transcription factor *SP1* were upregulated in IDA (Figure 3a).

**Fig 3.**
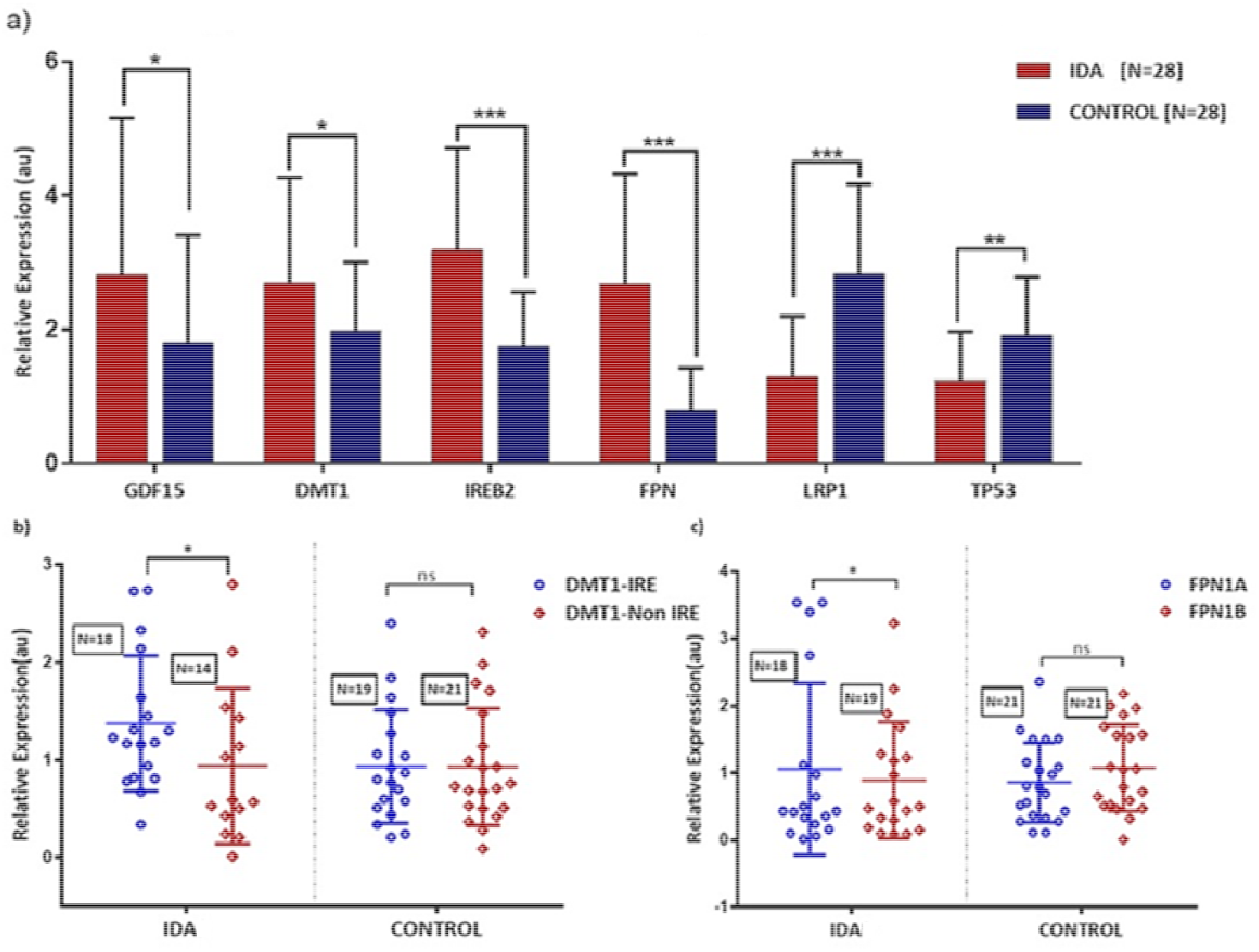
Figure 3 : a) Differentially expressed placental mRNA expression of iron traffickers in IDA and control group quantified using real time PCR. The expression level was normalized to β-actin. b&c) DMT1 and FPN isoforms mRNA expressions compared within IDAP and control groups. Data are presented as mean ± SD. N=28 in each group. Statistical significance was calculated using Student’s t-test (two-tailed t-test) and the P-values are denoted as NS, not significant, *P=0.05, **P=0.001 and ***P=0.0001.

Tumor suppressor gene TP53 suggested to which participates in maintenance of intracellular iron pool ^17^ and heme scavenger LRP1 were downregulated in IDA as compared to controls. No significant association was observed between mRNA expressions of above-mentioned iron transporters with respective downstream protein expression.

In IDA, placental TFRC did not differ at mRNA level. Under low iron levels (IDA), increased *DMT1* mRNA expression was observed. We found a positive correlation between *IREB2* mRNA expression and maternal serum ferritin (r=0.519, p=0.027). Upregulated cellular *FPN1* mRNA levels were associated with increased expression of *IREB2* mRNA (r=0.635, p=0.005).

At the protein level, GDF15, DMT1, TFRC and FPN1 were abundantly present in placenta (Figure 4).

**Fig 4:**
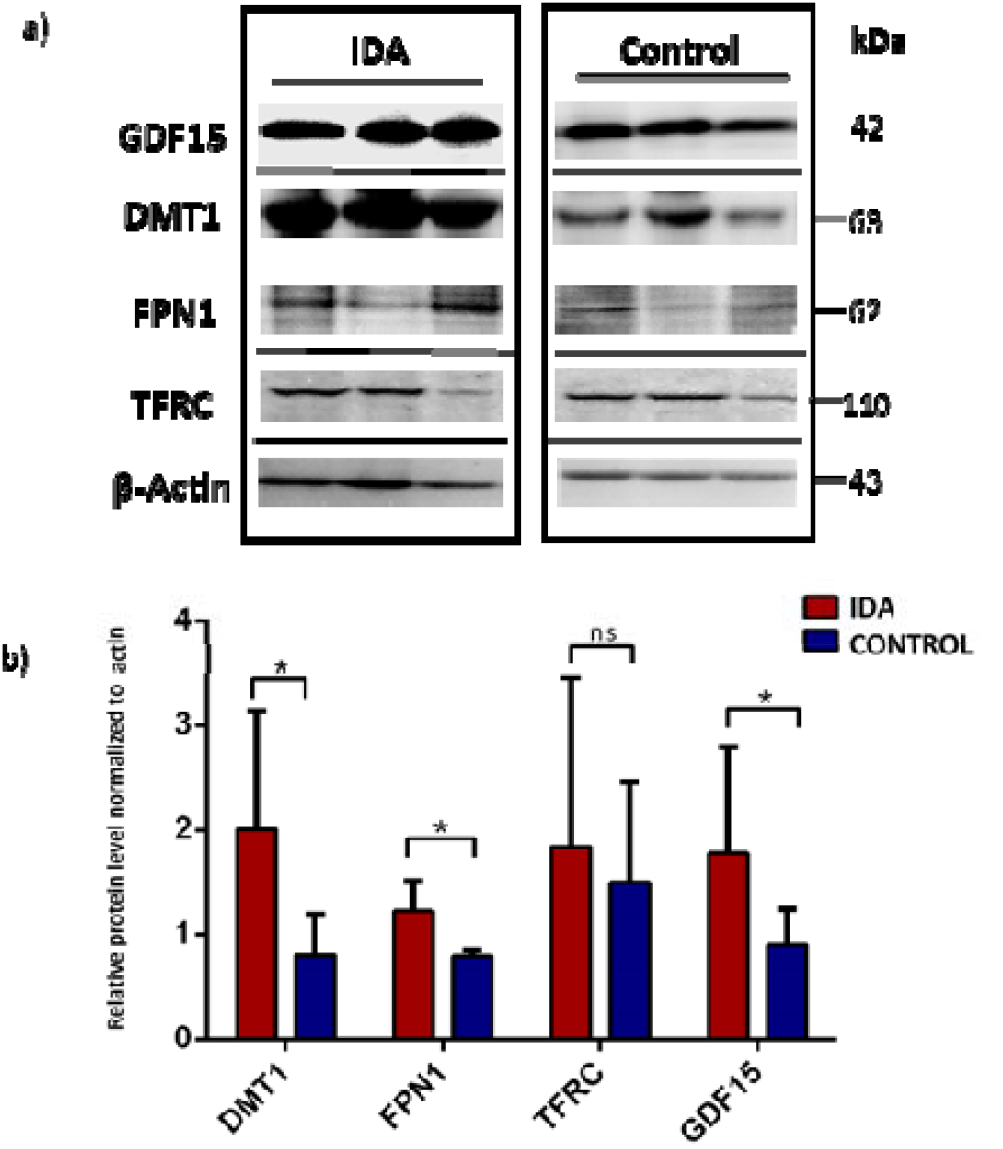
Western blot demonstration of placental protein expression of iron traffickers in IDA and control group.

FPN1 protein showed a trend towards significant association with maternal Hb levels (r=0.583; p=0.060). DMT1, FPN1 and GDF15 protein was significantly increased in IDA respectively (p=0.019, p=0.051, p=0.033). Positive associations were evident between placental GDF15, DMT1 and TFRC proteins in IDA.

Maternal Fe status indicators had no association with placental iron transporters at the mRNA level (Supplementary Table 3). There were no significant associations between *DMT1* protein expression with maternal and fetal iron status indicators (Supplementary Table 3).

On the other hand, heme uptake mediated by placental heme receptor LRP1 was differentially expressed in IDA (p=0.004) as compared to controls. LRP1 mRNA expression was significantly influenced by maternal ferritin levels (r=0.470; p=0.049), indicating the placental heme utilisation supports fetal iron demands. However, the heme scavenger CD163 and exporter FLVCR1 were not differentially expressed.

Several studies in line have evidenced the role of TP53 in maintenance of iron homeostasis, where it has been shown to influence hepcidin and ferritin levels ^17^. Zhang et al., have observed that loss of TP53 levels in iron overload mice had elevated serum iron levels. HAMP promoter region has a p53 putative responsive element which could be activated by P53. In our study, we observed placental TP53 mRNA expression significantly elevated in the iron deficient group. Interestingly, the placental TP53 mRNA expression had positive association with placental *GDF15* mRNA and maternal hepcidin concentration (r=0.642, p=0.004; r=0.492, p=0.038). However, this relation needs to be further studied.

Gestational age (39±1.2 weeks) had positive influence on placental iron traffickers including TFRC, LRP1 in IDA (r=0.568, p=0.017; r=0.625, p=0.007). Expression of Iron transport molecules in placenta did not influence neonatal birth weight. Maternal and fetal haemoglobin levels were not associated with the expression of placental iron transporters.

In control group, significant observation was a negative correlation between placental GDF15 mRNA and fetal Hb (r=-0.446, p=0.022).

### 3.5 Fetal iron transport by placental iron traffickers

Fetal ferritin was related to protein abundance of GDF15 (β=0.516 ;P=0.050)and Ferroportin (β=0.719 ;P<0.019). Fetal hepcidin-ferritin ratio had association with placental *FPN1* mRNA(β=0.532 ;P=0.028). These results indicate that fetal iron status regulates placental iron traffickers for iron transport towards fetus.

### 3.6 Splice variants in IDA placental iron transport

All spice variant transcripts of targeted iron transporters detected in the placental tissue were analysed. Alternative Splice variants of DMT1, FPN1, TFRC, SP1 and SLC46A1 were qualitatively confirmed. We observed differentially expressed isoforms of DMT1 and FPN1 in IDA. *FPN1* mRNA isoforms had increased expression in iron deficient cohort. *FPN1A* with 5’-Iron Regulatory Element (IRE) [FPN1A] expression was increased in IDA as compared to *FPN1B. DMT1A* mRNA isoform containing IRE was stabilized under iron deficient condition in IDA (p=0.05) (Fig 2b). *DMT1A* was positively associated with *IREB2* (r=0.512, p=0.018) and *FPN1A* (r=0.625, p=0.006). SP1 responsible for transcriptional response to iron deprivation had significant association with increased expression of *FPN 1A* (R²=0.478; p=0.003) and *FPN 1B* (R²=0.625; p=0.006).

## 4 Discussions

Iron deficiency anaemia in pregnancy is the most common public health concern affecting around 80% of pregnant women worldwide^18^. In South and South East Asian (SSEA) countries, prevalence of maternal anemia is estimated around 52%^19^. In India, 53% pregnant women have iron deficiency anemia^20^. Here, we investigated how maternal and fetal iron status relates to placental iron transporters’ expression.

In our study, despite iron supplementation, 22% pregnant women between 20-35 years old had iron deficiency anaemia at delivery. A similar finding was observed in Turkish pregnant women (18.7%)^14^. Iron deficiency also occurred in Gambian pregnant women regardless of iron supplementation^21^.

Most of them were mildly anaemic(Mean Hb – 9.2±0.66g/dl) identical to a study by Tabrizi et al., in Iranian pregnant women (Mean Hb level of 8.99±0.80g/dl)^15^. The risk factors such as maternal age, gestational age at delivery, gravida and consanguinity had no significant effects on IDA. Conversely, other findings suggested association between age and anemia^22^. Association between maternal anemia and low birth weight of newborns has been documented^23^. However, we did not observe such associations in our study. Cord blood ferritin and hepcidin are common biomarkers used to determine neonatal iron status at birth^24^. In our study, neonates born to iron deficient mothers had normal cord blood ferritin levels, thus confirming normal iron status in neonates.

Hepcidin, a systemic iron regulatory hormone, was suppressed in iron deficient cohort when compared to controls as observed earlier^25^. Maternal hepcidin had no association either with maternal or fetal iron status in IDA group. The decline in iron stores and hepcidin concentration at term pregnancy was also reported in Finland pregnant women^26^. Maternal hepcidin and iron status had no association with fetal iron regulators in IDA as reported in several studies, indicating the independent regulation of fetal iron status and fetal hepcidin^26^.

Several molecules are involved in iron trafficking between mother and foetus, whose regulation is still not clearly understood. Here we show that maternal iron deficiency did not affect fetal iron status; rather it had association with placental iron traffickers. During iron deficiency, TFRC was not differentially expressed at the transcriptional level, but its protein expression was increased in IDA. This finding was consistent with Sangkhae et al., human pregnancy model^24^. Iron transporters *DMT1* and *FPN1* mRNA expression were significantly elevated in IDA. This result signifies that maternal iron deficiency induces placental iron towards foetal circulation. And this also correlated with the increased expression of cellular iron-regulatory protein *IREB2*. Hence maternal anaemia has impact on placental iron traffickers for increased iron for foetal usage.

GDF15, an anti-inflammatory cytokine belongs to TGFβ superfamily, is highly expressed during pregnancy in the second and third trimester^27^,^16^. Decreased levels of serum GDF15 was reported in preeclampsia and miscarriage^27^,^9^. In accordance with other studies, we found augmented GDF15 concentration in pregnant women^9^. GDF15 suppresses hepcidin in β-thalassemia^28^ and CDA^29^, may control hepcidin in pregnancy. We did not find association of hepcidin and ferritin with GDF15 levels in both mother and fetus. However, we observed strong expression of GDF15 in placenta at mRNA and protein level. GDF15 protein expression had positive associations with TFRC, DMT1, SP1 and TP53 proteins reflecting essential role of GDF15 in placental iron regulation. This is supported by the fact that transcription factors SP1 and TP53 are involved in GDF15 upregulation in erythroid cells^16^. Based on this analysis we suggest SP1 and TP53 increase GDF15 expression in placenta.

We postulate that fetal iron status may regulate placental GDF15 and ferroportin for adequate transfer of iron towards fetal circulation as observed by their positive correlations. However, the function of placental GDF15 in iron regulation needs to be further characterised.

Alternative splicing of pre-mRNA produces multiple mRNA transcripts from a gene through post transcriptional mechanism^30^. This is the first study, to best of our knowledge to measure isoforms of multiple non-heme iron transport proteins except FPN1 in human placental tissue in iron deficient and iron replete groups.

Invitro studies have observed DMT1A expression in duodenal enterocytes and DMT1B in leukocytes^31^. In our study, DMT1A isoform with IRE is significantly increased in iron deficiency helping in increased iron absorption. FPN1 isoforms localisation in erythroid cells were initially reported by Cianetti et al^32^. Increased expression of FPN 1A in placenta mirrors the recent data of Sangkhae and colleagues^24^.

In controls, even under replete maternal iron stores, reduced hepcidin levels allowed iron mobilisation from stores into maternal circulation. Similar to several authors^25^,^33^,^34^, we also have observed positive association between hepcidin and ferritin in healthy pregnant women and no correlation between maternal and fetal iron status.

This is one of the few studies to be carried out in humans to understand how iron status is maintained in the foetus even when the mother has depleted iron stores and anaemia. Several interesting findings have been identified; however, this study is limited by numbers. It was not possible to do radio iron transfer in the iron deficient pregnant women. Some correlations could have thrown more light with functional studies like EMSA.

The present observations in the group of iron deficient pregnant women demonstrated that lower iron status and hepcidin levels induce increased iron mobilisation from iron stores. Foetal iron status was independent of maternal ferritin and hepcidin levels. Figure 5 summarizes the proposed mechanism of iron regulation in IDA.

**Fig 5:**
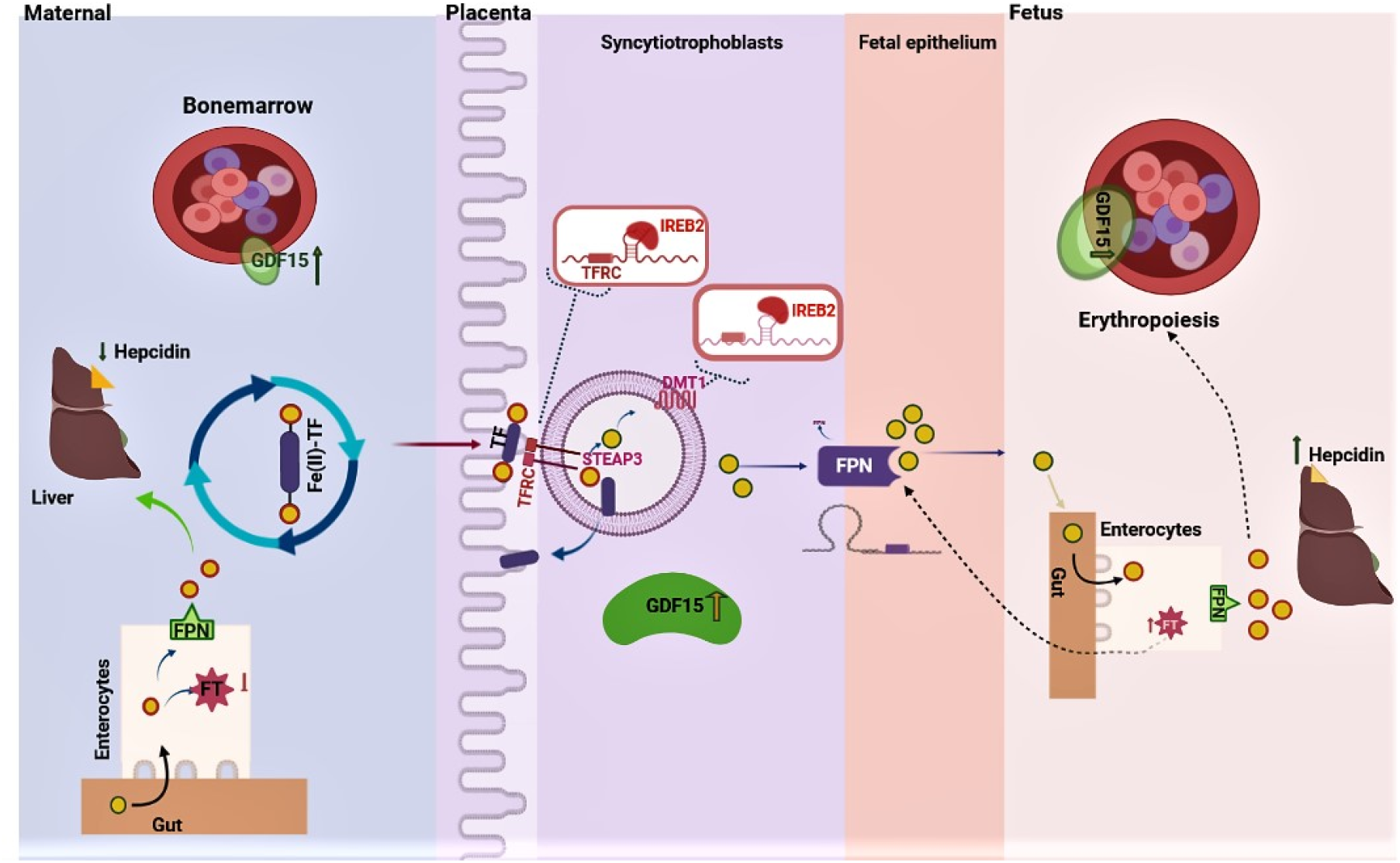
Proposed mechanism of iron regulation in maternal-placenta-fetal pathway in Iron deficiency anaemia of pregnancy

Under maternal iron deficiency, Fe absorbed in duodenum are partially stored in ferritin (FT) reservoir and maximum Fe released into the circulation via ferroportin (FPN). In the circulation, Fe is transported in a complex with transferrin (TF) to hepatocytes and bone marrow. Hepcidin suppression elevates FPN expression and results in maximum Fe absorption by mobilising Fe from internal stores. Increased erythropoiesis in pregnancy induces increased GDF15 production. From maternal circulation, TF-Fe (II) complex binds to transferrin receptor (TFRC) in apical side of syncytiotrophoblasts and gets endocytosed. Acidified vesicle allows oxidation of ferrous into ferric Fe via STEAP3 and exported into cytoplasm through DMT1. IRP2 activity on TFRC and DMT1 promotes their transcription, resulting in increased iron transport. Abundant expression of GDF15 in placenta might also involve in regulation of Fe transport. From basal side of syncytiotrophoblasts, Fe is exported via FPN into the fetal circulation. In fetus, internal iron stores regulate placental FPN and GDF15, thereby increasing iron endowment and maintains fetal iron homeostasis.

In maternal iron deficiency, placental iron regulators were upregulated and had association with fetal iron status, which implies that placenta allows excess iron transport to fetus at the expense of mother.

## Authorship

Contribution: S.S. performed the experiments, analysed the data, prepared tables, figures and wrote the paper; E.S. and S.S. performed the statistical analysis; A.G.C. and V.J.A. recruited and treated the pregnant women. R.A., A.G.C., B.G., provided critical advice and edited the paper. P.L gave valuable inputs to the study and reviewed the manuscript. E.S. conceptualized and supervised the research, organised the data, edited the paper. All authors approved read and approved the final manuscript.

## Acknowledgement

We thank Dr. Arun Jose and Dr. Joseph Bondu for helping in biochemical analysis.

Correspondence: Eunice Sindhuvi, Department of Hematology, Christian Medical College, Vellore-632004,TamilNadu,India

## Ethics approval

The study was approved by Institutional Review Board (Ethics committee) of Christian Medical College (CMC) at Vellore, India, (IRB No.9360 and dated 25-03-2015).

## Consent to participate

Informed consent was obtained from all individual participants included in the study.

## References

1. Delaney KM, Guillet R, Fleming RE, et al. Umbilical Cord Serum Ferritin Concentration is Inversely Associated with Umbilical Cord Hemoglobin in Neonates Born to Adolescents Carrying Singletons and Women Carrying Multiples. The Journal of Nutrition. 2019;149(3):406–415. doi:10.1093/jn/nxy286

2. Rahman KM, Ali KM, Vijayalakshmi S, Ramkumar S, Hashmi G. Prevalence of Iron Deficiency Anaemia and its Associated Factors among Reproductive Age Women in a Rural Area of Karaikal, Puducherry, India. JCDR. Published online 2019. doi:10.7860/JCDR/2019/36623.12706

3. Kapil U, Bhadoria AS. National Iron-plus initiative guidelines for control of iron deficiency anaemia in India, 2013. Natl Med J India. 2014;27(1):27–29.

4. Bothwell TH. Iron requirements in pregnancy and strategies to meet them. The American Journal of Clinical Nutrition. 2000;72(1):257S–264S. doi:10.1093/ajcn/72.1.257S

5. Koenig MD, Tussing-Humphreys L, Day J, Cadwell B, Nemeth E. Hepcidin and Iron Homeostasis during Pregnancy. Nutrients. 2014;6(8):3062–3083. doi:10.3390/nu6083062

6. Sangkhae V, Nemeth E. Placental iron transport: The mechanism and regulatory circuits. Free Radic Biol Med. 2019;133:254–261. doi:10.1016/j.freeradbiomed.2018.07.001

7. Young MF, Griffin I, Pressman E, et al. Maternal Hepcidin Is Associated with Placental Transfer of Iron Derived from Dietary Heme and Nonheme Sources1234. J Nutr. 2012;142(1):33–39. doi:10.3945/jn.111.145961

8. Nicolas G, Bennoun M, Porteu A, et al. Severe iron deficiency anemia in transgenic mice expressing liver hepcidin. Proc Natl Acad Sci U S A. 2002;99(7):4596–4601. doi:10.1073/pnas.072632499

9. Finkenstedt A, Widschwendter A, Brasse-Lagnel CG, et al. Hepcidin is correlated to soluble hemojuvelin but not to increased GDF15 during pregnancy. Blood Cells, Molecules, and Diseases. 2012;48(4):233–237. doi:10.1016/j.bcmd.2012.02.001

10. Maltepe E, Fisher SJ. Placenta: The Forgotten Organ. Annu Rev Cell Dev Biol. 2015;31(1):523–552. doi:10.1146/annurev-cellbio-100814-125620

11. Bradley J, Leibold EA, Harris ZL, et al. Influence of gestational age and fetal iron status on IRP activity and iron transporter protein expression in third-trimester human placenta. Am J Physiol Regul Integr Comp Physiol. 2004;287(4):R894–901. doi:10.1152/ajpregu.00525.2003

12. Chong WS, Kwan PC, Chan LY, Chiu PY, Cheung TK, Lau TK. Expression of divalent metal transporter 1 (DMT1) isoforms in first trimester human placenta and embryonic tissues. Hum Reprod. 2005;20(12):3532–3538. doi:10.1093/humrep/dei246

13. Sangkhae V, Fisher AL, Wong S, et al. Effects of maternal iron status on placental and fetal iron homeostasis. The Journal of Clinical Investigation. Published online October 29, 2019. doi:10.1172/jci127341

14. Kavak E, Kavak salih burcin. The association between anemia prevalence, maternal age and parity in term pregnancies in our city. Perinatal Journal. 2017;25:6–10. doi:10.2399/prn.17.0251002

15. Moghaddam Tabrizi F, Barjasteh S. Maternal Hemoglobin Levels during Pregnancy and their Association with Birth Weight of Neonates. Iran J Ped Hematol Oncol. 2015;5(4):211–217.

16. Tanno T, Noel P, Miller JL. Growth differentiation factor 15 in erythroid health and disease. Curr Opin Hematol. 2010;17(3):184–190. doi:10.1097/MOH.0b013e328337b52f

17. Zhang J, Chen X. p53 tumor suppressor and iron homeostasis. The FEBS Journal. 2019;286(4):620–629. doi:https://doi.org/10.1111/febs.14638

18. WHO | Intermittent iron and folic acid supplementation during pregnancy in malaria-endemic areas. WHO. Accessed January 4, 2021. http://www.who.int/elena/titles/intermittent_iron_pregnancy_malaria/en/

19. Sunuwar DR, Singh DR, Chaudhary NK, Pradhan PMS, Rai P, Tiwari K. Prevalence and factors associated with anemia among women of reproductive age in seven South and Southeast Asian countries: Evidence from nationally representative surveys. Cardoso MA, ed. PLoS ONE. 2020;15(8):e0236449. doi:10.1371/journal.pone.0236449

20. National Family Health Survey. Accessed January 20, 2021. http://rchiips.org/nfhs/

21. Bah A, Pasricha SR, Jallow MW, et al. Serum Hepcidin Concentrations Decline during Pregnancy and May Identify Iron Deficiency: Analysis of a Longitudinal Pregnancy Cohort in The Gambia. The Journal of Nutrition. 2017;147(6):1131–1137. doi:10.3945/jn.116.245373

22. Gautam S, Min H, Kim H, Jeong HS. Determining factors for the prevalence of anemia in women of reproductive age in Nepal: Evidence from recent national survey data. PLOS ONE. 2019;14(6):e0218288. doi:10.1371/journal.pone.0218288

23. Widiyanto J, Lismawati G. Maternal age and anemia are risk factors of low birthweight of newborn. Enferm Clin. 2019;29:94–97. doi:10.1016/j.enfcli.2018.11.010

24. Sangkhae V, Fisher AL, Wong S, et al. Effects of maternal iron status on placental and fetal iron homeostasis. J Clin Invest. 2020;130(2):625–640. doi:10.1172/JCI127341

25. Kulik-Rechberger B, Kościesza A, Szponar E, Domosud J. Hepcidin and iron status in pregnant women and full-term newborns in first days of life. Ginekologia Polska. 2016;87(4):288–292. doi:10.17772/gp/62202

26. Rehu M, Punnonen K, Ostland V, et al. Maternal serum hepcidin is low at term and independent of cord blood iron status. European Journal of Haematology. 2010;85(4):345–352. doi:https://doi.org/10.1111/j.1600-0609.2010.01479.x

27. Chen Q, Wang Y, Zhao M, Hyett J, da Silva Costa F, Nie G. Serum levels of GDF15 are reduced in preeclampsia and the reduction is more profound in late-onset than earlyonset cases. Cytokine. 2016;83:226–230. doi:10.1016/j.cyto.2016.05.002

28. Tanno T, Bhanu NV, Oneal PA, et al. High levels of GDF15 in thalassemia suppress expression of the iron regulatory protein hepcidin. Nat Med. 2007;13(9):1096–1101. doi:10.1038/nm1629

29. Tamary H, Shalev H, Perez-Avraham G, et al. Elevated growth differentiation factor 15 expression in patients with congenital dyserythropoietic anemia type I. Blood. 2008;112(13):5241–5244. doi:10.1182/blood-2008-06-165738

30. Black A, Gamarra J, Giudice J. More than a messenger: Alternative splicing as a therapeutic target. Biochim Biophys Acta Gene Regul Mech. 2019;1862(11-12):194395. doi:10.1016/j.bbagrm.2019.06.006

31. Tabuchi M, Tanaka N, Nishida-Kitayama J, Ohno H, Kishi F. Alternative Splicing Regulates the Subcellular Localization of Divalent Metal Transporter 1 Isoforms. Mol Biol Cell. 2002;13(12):4371–4387. doi:10.1091/mbc.E02-03-0165

32. Cianetti L, Segnalini P, Calzolari A, et al. Expression of alternative transcripts of ferroportin-1 during human erythroid differentiation. Haematologica. 2005;90(12):1595–1606.

33. van Santen S, Kroot JJC, Zijderveld G, Wiegerinck ET, Spaanderman MEA, Swinkels DW. The iron regulatory hormone hepcidin is decreased in pregnancy: a prospective longitudinal study. Clin Chem Lab Med. 2013;51(7):1395–1401. doi:10.1515/cclm-2012-0576

34. Lorenz L, Herbst J, Engel C, et al. Gestational Age-Specific Reference Ranges of Hepcidin in Cord Blood. NEO. 2014;106(2):133–139. doi:10.1159/000360072

