## Supplementary data for "Impact of maternal iron deficiency anaemia on fetal iron status and placental iron transporters in human pregnancy"

**Supplemental Table 1: List of genes included in custom array panel**

| **Sno** | **Gene Symbol** |
|---|---|
| 1 | GDF15 |
| 2 | SLC11A2 |
| 3 | SLC40A1 |
| 4 | HEPHL1 |
| 5 | FTL |
| 6 | FLVCR1 |
| 7 | HIF1A |
| 8 | TP53 |
| 9 | ACO1 |
| 10 | IREB2 |
| 11 | PGF |
| 12 | SP1 |
| 13 | SLC46A1 |
| 14 | HFE |
| 15 | CD163 |
| 16 | LRP1 |
| 17 | TFR2 |
| 18 | TWSG1 |
| 19 | TFRC |
| 20 | GAPDH |
| 21 | ACTB |

**Supplemental Table 2: Primary and Secondary antibodies used in western blot**

| Target Protein | Primary antibody | Dilution |
|---|---|---|
| FPN1 | Rabbit polyclonal antibody (NBP1-21502,Novus biologicals) | 1:1000 |
| DMT1 | Rabbit polyclonal antibody (Cat no: A10231,Abclonal) | 1:1000 |
| TFR1 | Rabbit polyclonal antibody (Cat no: A18083, Abclonal) | 1:1000 |
| GDF15 | Mouse monoclonal antibody (sc-515675, Santa cruz biotechnology) | 1:1000 |
| β-Actin | Mouse monoclonal antibody (Sc-47778, Santa cruz biotechnology) | 1:1000 |

| Secondary antibody |
|---|
| Anti-mouse IgG HRP (Santa cruz biotechnology, sc-2005) |
| Anti-rabbit IgG HRP (Cell Signaling #7074) |
| Anti-goat IgG HRP(Santa cruz biotechnology, sc-2020) |

**Supplemental Table 3: Correlations of mRNA expressions and protein abundance of placental Fe transporters with maternal and cord blood Fe status indicators.**

| | Mother | | | | Cordblood | | | |
|---|---|---|---|---|---|---|---|---|
| Placental Fe transporters | Hb | Ferritin | Hepcidin | GDF15 | Hb | Ferritin | Hepcidin | GDF15 |
| mRNA DMT1 | 0.722 | 0.091 | 0.541 | 0.893 | 0.820 | 0.775 | 0.414 | 0.854 |
| mRNA TFRC | 0.912 | 0.101 | 0.486 | 0.958 | 0.766 | 0.852 | 0.151 | 0.405 |
| mRNA FPN | 0.698 | 0.250 | 0.772 | 0.723 | 0.645 | 0.468 | 0.937 | 0.854 |
| mRNA GDF15 | 0.977 | 0.210 | 0.747 | 0.958 | 0.763 | 0.480 | 0.268 | 0.778 |
| mRNA IREB2 | 0.797 | 0.027 | 0.580 | 0.657 | 0.868 | 0.801 | 0.918 | 0.499 |
| Protein DMT1 | 0.679 | 0.734 | 0.158 | 0.986 | 0.541 | 0.587 | 0.217 | 0.853 |
| ProteinTFRC | 0.609 | 0.824 | 0.703 | 0.761 | 0.602 | 0.725 | 0.736 | 0.642 |
| Protein FPN | 0.060 | 0.450 | 0.958 | 0.417 | 0.769 | 0.048 | 0.626 | 0.244 |
| Protein GDF15 | 0.479 | 0.155 | 0.252 | 0.589 | 0.687 | 0.017 | 0.193 | 0.128 |

**Supplemental Table 3:** Correlation matrix of placental mRNA and proteins with maternal and fetal iron parameters are tabulated. Significantly correlated values highlighted in blue.

**Supplemental Figure 1**

Supplemental Fig 1: VNTR analysis of Maternal and cord blood and placental tissue DNA (N=2)

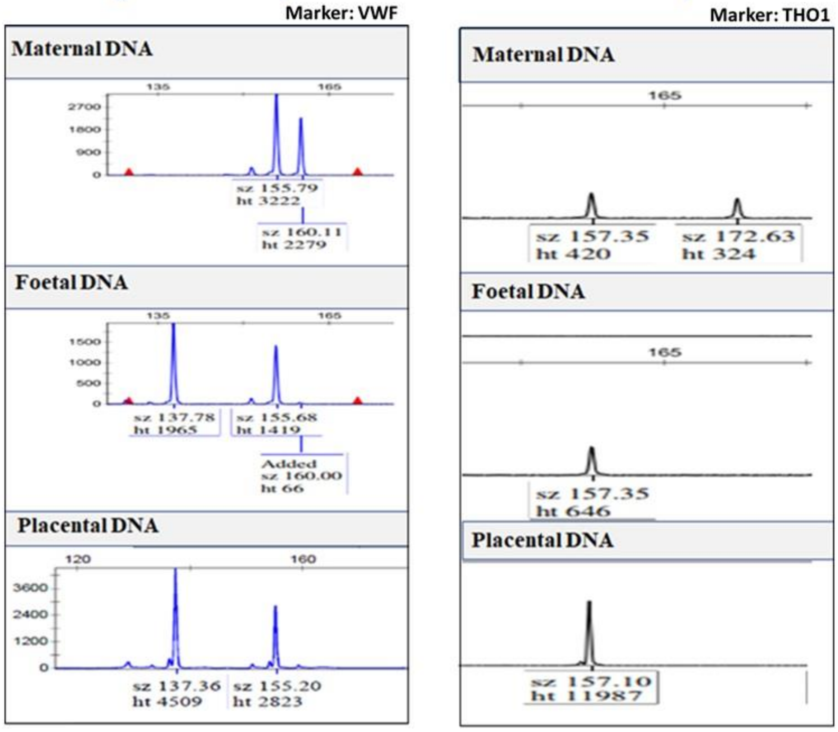

**Supplemental Figure 2**

**Study Participants**

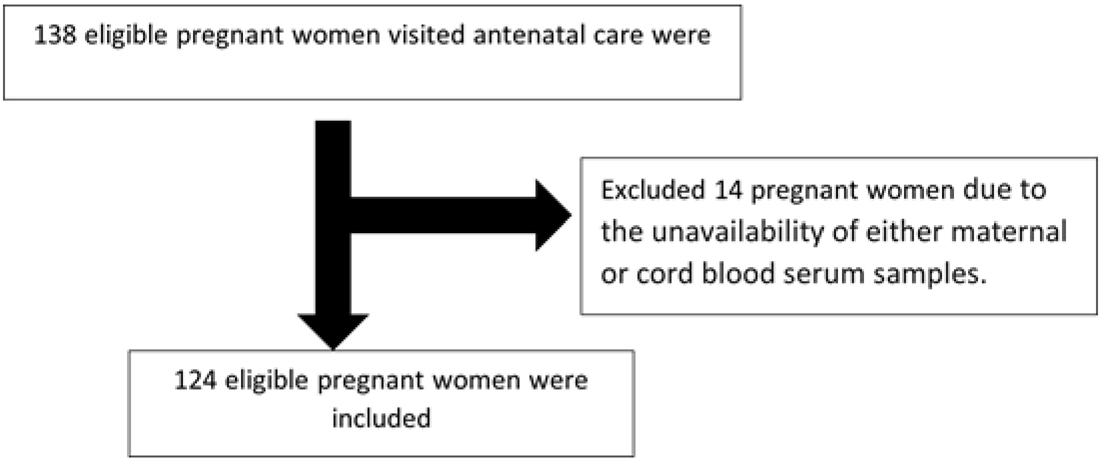
